# The role of location propagation for translocon complex relocation during eukaryogenesis

**DOI:** 10.1101/2023.10.04.560972

**Authors:** Isaac Carilo, Yosuke Senju, Robert C. Robinson

**Affiliations:** Research Institute for Interdisciplinary Science, Okayama University, Okayama 700-8530, Japan; School of Biomolecular Science and Engineering (BSE), Vidyasirimedhi Institute of Science and Technology (VISTEC), Rayong, 21210, Thailand

## Abstract

In all domains of life, the translationally-active ribosome-translocon complex inserts nascent transmembrane proteins into, and processes and transports signal peptide-containing proteins across, membranes^1,2^. Eukaryotic translocons are anchored in the endoplasmic reticulum, while the prokaryotic complexes reside in cell membranes. DNA sequence analyses indicate that the eukaryotic Sec61/OST/TRAP translocon is inherited from an Asgard archaea ancestor^3,4^. However, the mechanism for translocon migration from a peripheral membrane to an internal cellular compartment (the proto-endoplasmic reticulum) during eukaryogenesis is unknown. Here we show that Asgard and eukaryotic ribosome-translocon complexes are intercompatible. We find that fluorescently-tagged Asgard translocon proteins from *Candidatus* Prometheoarchaeum syntrophicum strain MK-D1^5^, a Lokiarchaeon confirmed to contain no internal cellular membranes, are targeted to the eukaryotic endoplasmic reticulum on ectopic expression. Our data demonstrate that the location of existing ribosome-translocon complexes, at the protein level, determines the future placement of yet to be translated translocon subunits. This principle predicts that during eukaryogenesis, under positive selection pressure, the relocation of a few translocon complexes to the proto-endoplasmic reticulum will have propagated the new translocon location, leading to their loss from the cell membrane.

---

There are many hypotheses for the origin of eukaryotic internal membranes^5–10^. However, it is challenging to reproduce the actual events during eukaryogenesis, which occurred more than 1.8 billion years ago (BYA)^11^, or to design meaningful experiments that can distinguish between these theories. Possibly, the best tools available in understanding the emergence of eukaryotic internal membranes are the transmembrane proteins associated with each membrane. In particular, the eukaryotic Sec61/OST/TRAP translocon resides in the endoplasmic reticulum (ER) membrane and mediates the insertion and N-glycosylation of transmembrane proteins that are being actively translated by ribosomes. By contrast, in prokaryotes the homologous SecYEG translocon sits in the cell membrane. These diametrically opposed locations provide an opportunity to probe how the Sec61/OST/TRAP translocon may have become relocalized during eukaryogenesis. Phylogenetic analyses of the proprotein translocase channel SecY/Sec61 and the oligosaccharide-transfer (OST) complex catalytic subunit (STT3) indicate that the eukaryotic versions of these core translocon subunits are more closely related to their Asgard archaea counterparts than to other prokaryotic homologs^4,12^. Asgard archaea are predicted to have a complete Sec61αβγ complex and many of the components of the OST and translocon-associated proteins (TRAP) complexes^3^. In particular, sequence searches predict genes for the Sec61αβγ complex, OST1 (ribophorin), OST3/6 and STT3, and TRAP subunits α, β and γ in MK-D1.

In order to probe the similarities between the MK-D1 and eukaryotic proprotein signal sequences, we ran the MK-D1 genome-predicted protein sequences in the signal sequence prediction software, SignalP 6.0^13^. Many MK-D1 transmembrane proteins were predicted to have eukaryotic-like signal sequences, including OST1, TRAPα and TRAPβ, with probabilities of 0.77, 1.00 and 0.99, respectively, this compares with the probabilities for the human proteins of 1.00, 0.81 and 1.00, respectively (Fig. 1). However, the eukaryotic subcellular localization software DeepLoc 2.0^14^ predicted a variety of eukaryotic membrane localizations for these MK-D1 membrane proteins, in comparison to the strong ER localization predicted for the human translocon subunits (Fig. 1), in line with the evolution of complex membrane protein processing within eukaryotic cells. SignalP 6.0 did not predict signal peptides for the homologs of eukaryotic ER chaperones^15^ in MK-D1, or in any other Asgard organism, indicating that the Asgard chaperone homologs are cytoplasmic.

**Figure 1.**
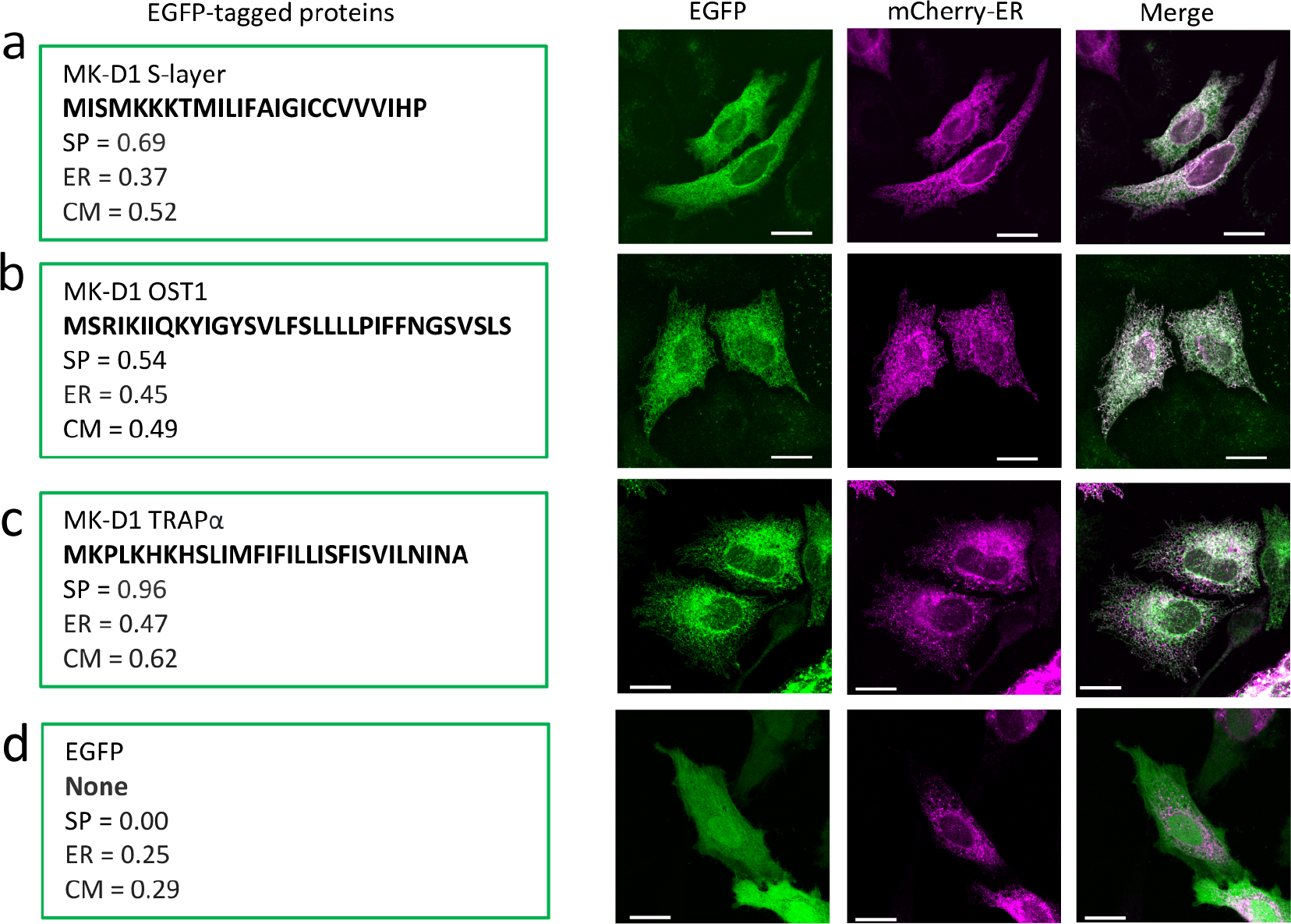
Localization of MK-D1 full length signal peptide-containing proteins on transfection in HeLa cells. Cells were co-transfected to express EGFP-fused signal peptide-containing proteins and an ER-localizing mCherry construct. At 24 h post-transfection, cells were fixed and imaged using the confocal microscope. The EGFP (green), mCherry (purple) and merged images are shown. **a**, MK-D1 S-layer protein. **b**, MK-D1 OST1. **c**, MK-D1 TRAPα. **d**, EGFP alone. The signal peptide sequences for each MK-D1 protein (bold), SignalP 6.0 signal peptide prediction scores (SP), and DeepLoc 2.0 localization probabilities for the endoplasmic reticulum (ER) and cell membrane (CM) are given for each EGFP construct. Scale bar = 20 μm.

To experimentally determine whether MK-D1 proproteins exhibit a preferred location in eukaryotic cells, we ectopically expressed MK-D1 OST1, TRAPα and the cell surface S-layer protein as EGFP fusion proteins in HeLa cells (Fig. 1a-c). In all three cases, these MK-D1 cell surface proteins co-localized with an mCherry ER marker, and did not localize to the cell membrane. By contrast, EGFP alone showed no co-localization with the ER marker (Fig. 1d). These data indicate that heterologously expressed cell surface Asgard proproteins are translated and processed at the ER, where the eukaryotic translocase is located.

To demonstrate that MK-D1 signal peptides are responsible for ER localization, we expressed a series of signal sequences fused to EGFP in HeLa cells (Fig. 2). Signal peptides from the MK-D1 S-layer protein, TRAPα, TRAPγ and OST1 all localized to the ER, as did the positive control human OST1 (Fig. 2c) and did not show the diffuse expression pattern of EGFP alone (Fig. 1d). Thus, the MK-D1 signal peptides are responsible for directing localization to the ER. In order to further assess the compatibility of Asgard signal peptides with eukaryotic Sec61α, we used AlphaFold2 (AF2)^16,17^ to predict the complex structures of human Sec61α with the MK-D1 signal peptides (Fig. 2). In each prediction, the MK-D1 signal peptides occupy the central channel of human Sec61α in a similar orientation to the structure of a signal peptide-engaged Sec61 complex^18^, and observed in the control AF2 prediction, human OST1 signal peptide with human Sec61α (Fig. 2c). Taken together, these data indicate that Asgard signal peptides are compatible with the eukaryotic Sec61α translocase, and when heterologously expressed in eukaryotic cells, Asgard signal peptide containing proteins are co-translated and translocated at the site of eukaryotic Sec61 localization, the ER.

**Figure 2.**
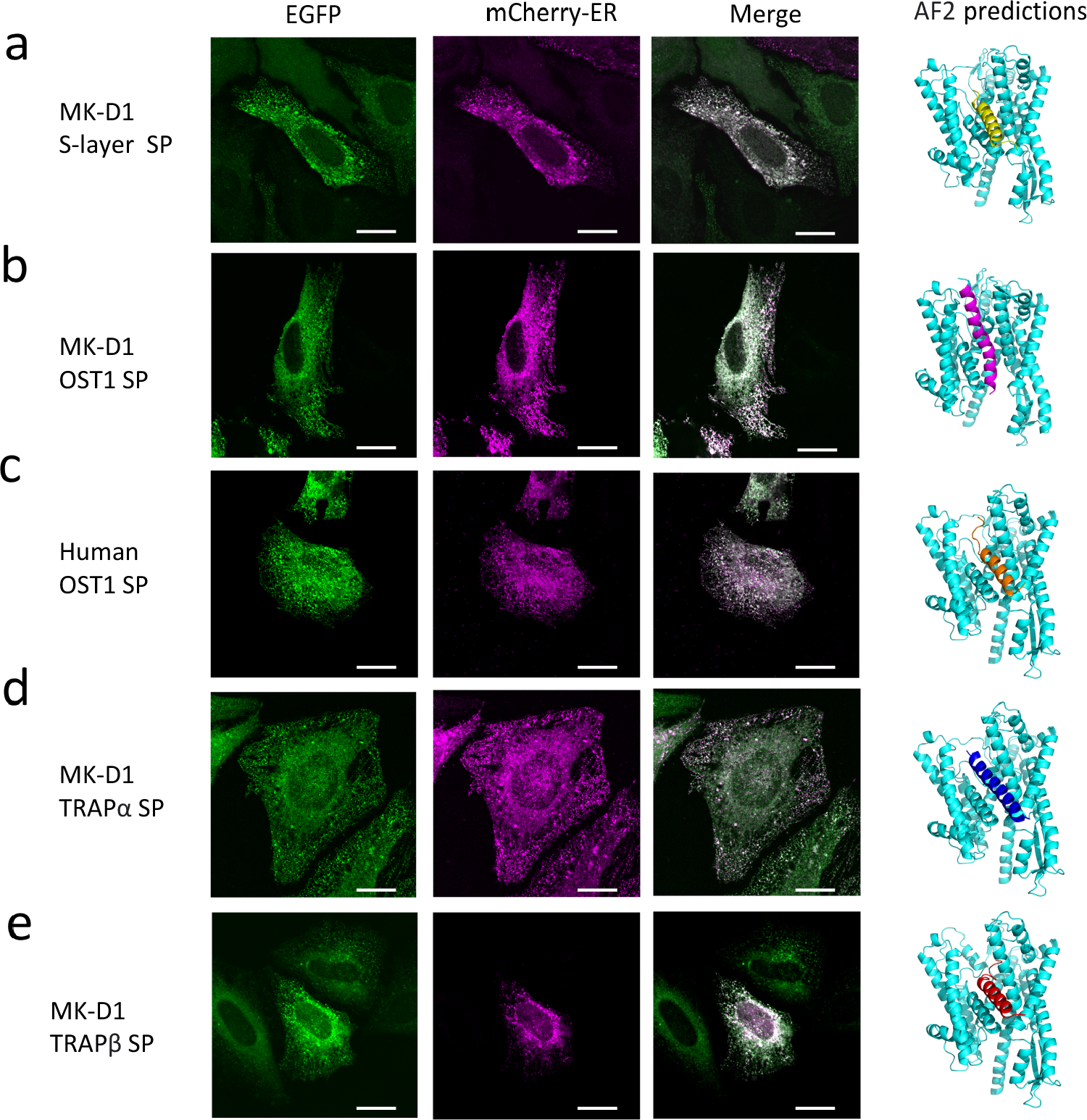
Localization of MK-D1 signal peptide-EGFP chimeras on transfection in HeLa cells. Signal peptides (SP) alone, from MK-D1 proteins, were fused to EGFP and co-transfected with the ER marker into HeLa cells as in Fig. 1. **a**, MK-D1 S-layer protein SP. **b**, MK-D1 OST1 SP. **c**, Human OST1 SP control. **d**, MK-D1 TRAPα SP. **e**, MK-D1 TRAPβ SP. AF2 co-predictions are shown as cartoons for the human Sec61α and each of the signal peptides. Sec61α, cyan; S-layer SP, yellow; MK-D1 OST1 SP, magenta; Human OST1 SP, orange; MK-D1 TRAPα SP, blue; MK-D1 TRAPβ SP, red. Scale bar = 20 μm.

To discover whether Asgard transmembrane proteins, which lack signal peptides, are preferentially located in eukaryotic cells, we expressed this class of proteins from the MK-D1 Sec61/OST/TRAP translocon as EGFP fusion proteins in HeLa cells. The three components of the MK-D1 Sec61αβγ proprotein translocase complex localized with the mCherry human Sec61αβγ subunits (Fig. 3), indicating that the MK-D1 Sec61 is directed to the same compartment as human Sec61, the ER. Finally, we co-expressed MK-D1 transmembrane proteins from the OST and TRAP complexes with the mCherry ER marker. EGFP tagged MK-D1 TRAPγ, OST3/6 and STT3 all colocalized with the ER marker (Fig. 4). Thus, the entire set of MK-D1 Sec61/OST/TRAP translocon subunits become located to the ER when expressed in eukaryotic cells. Taken together, these data establish the principle that the location of the an existing Sec61 complex, on engaging the ribosome, determines the site for the new Sec61/OST/TRAP translocons. Translocon location is inherited at a protein molecular level rather than at a genetic level, which has likely implications for ER maintenance, disease and our focus, biogenesis.

**Figure 3.**
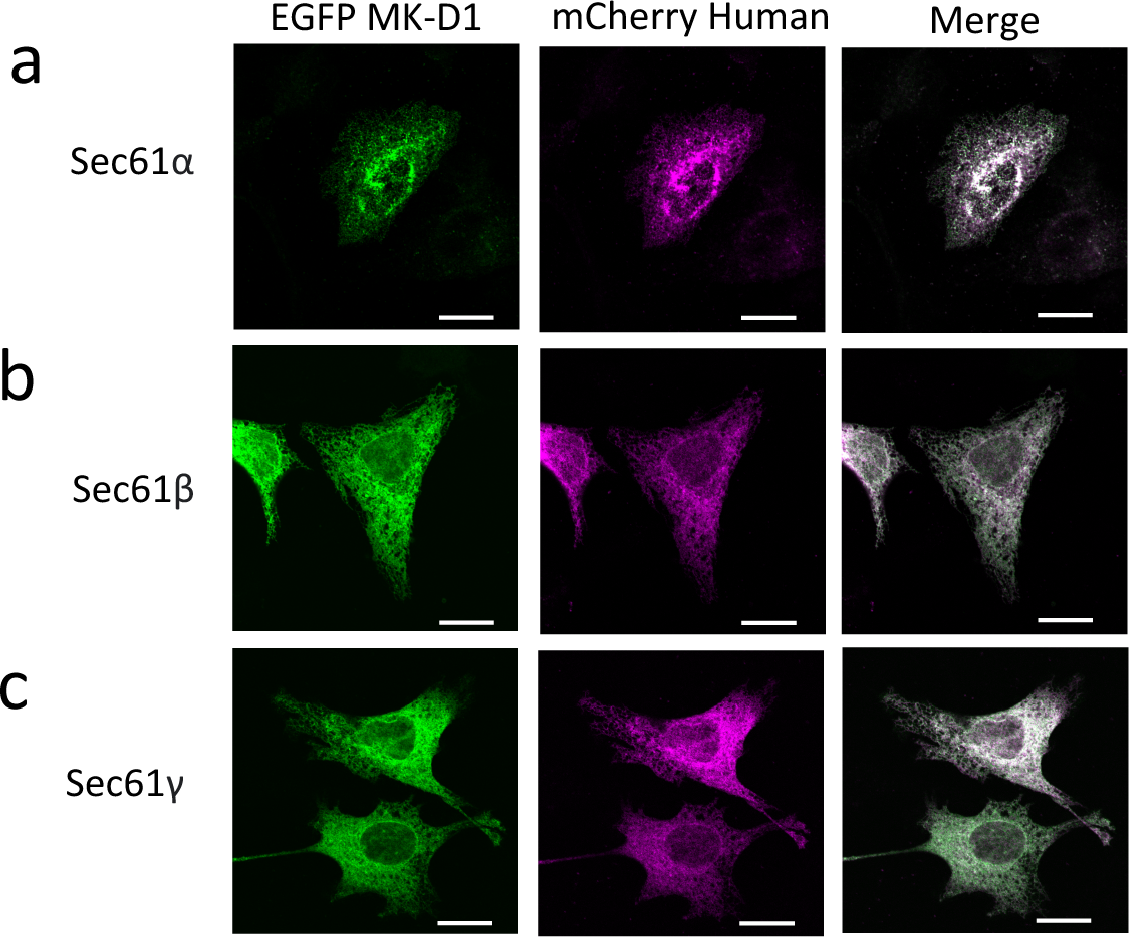
Localization of non-SP MK-D1 transmembrane Sec61 proteins on transfection in HeLa cells. **a-c**, Each of EGFP-tagged MK-D1 **a**, Sec61α, **b**, Sec61β and **c**, Sec61γ was co-transfected with its corresponding mCherry fused-human counterpart into HeLa cells. Scale bar = 20 μm.

**Figure 4.**
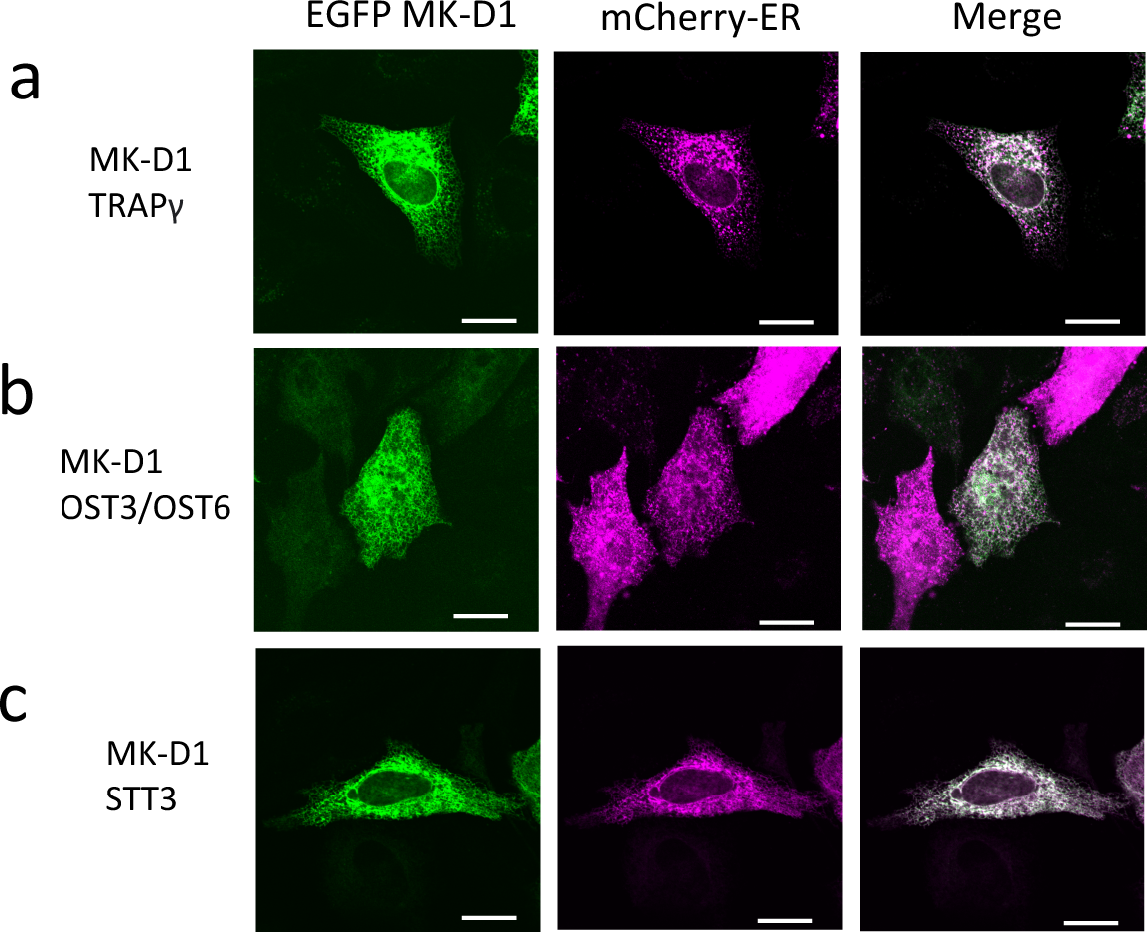
Localization of non-SP MK-D1 transmembrane TRAP and OST proteins on transfection in HeLa cells. **a-c**, HeLa cells were co-transfected to express each of EGFP-fused MK-D1 **a**, TRAPγ, **b**, OST3/OST6 and **c**, STT3 with the ER localizing mCherry construct. At 24 h post-transfection, cells were fixed and imaged with the confocal microscope. Scale bar = 20 μm.

The intercompatibility of Asgard and eukaryotic ribosome-translocon complexes has implications for ER biogenesis. These data are consistent with models of eukaryogenesis where the cell membrane is derived from an Asgard archaeon^20^. Furthermore, the proto-ER membrane should be derived from a membrane that allows docking of Asgard cytoplasmic ribosomes to the translocon in the cell membrane and proto-ER. Invagination or cell expansion models leading to the emergence of the proto-ER from the cell membrane are consistent with this scenario. However, such models require an additional step of the emergence of vesicle budding and fusion at the cell surface to complete the transport of transmembrane proteins from the proto-ER to cell membrane. Models of ER origin that involve membranes from endosymbionts residing within the Asgard archaeon have an initial problem in that the translocons are inverted and would not be able to engage Asgard cytoplasmic ribosomes. However, the emergence of early endocytic vesicle trafficking may have allowed for the shuttling of Asgard translocons, in the correct orientation, from the cell surface to the proto-ER.

The potential sources of proto-ER membrane require either the emergence of endocytic or exocytic vesicle transport. These processes are present in extant eukaryotes, and many homologs of eukaryotic-like vesicle transport proteins have been identified in Asgard archaea^3^, implying that the emergence of vesicle transport is a realistic step within an Asgard-derived cytoplasm. However, the lack of chaperones containing signal peptides in Asgard sequence databases suggests that a proto-ER, in which chaperone-assisted refolding occurs, is not present in the extant Asgard organisms sequenced to date. The mechanism described here, by which the existing translocon location directs future translocon distribution (translocon-location propagation) in the proto-eukaryote, creates an extended window for the proto-ER to evolve efficient protein modification and folding, and vesicle transport of transmembrane proteins. We suggest that once the quality of folded and modified transmembrane proteins from the proto-ER surpassed that made at the cell membrane, and the vesicle transport system became efficient, then translocon-location propagation would have relocated the entire population of translocons to the proto-ER.

## Acknowledgements

We thank the SPring-8 Synchrotron, Japan for facilities. This work was supported by the Moore-Simons Project on the Origin of the Eukaryotic Cell, grant number GBMF9743 (R.C.R.); Japan Society for the Promotion of Science (JSPS), grant number JP20H00476 (R.C.R.), JP19K23727 (Y.S.), JP23K05718 (Y.S.), JP23H04423 (Y.S.); and by JST CREST, grant number JPMJCR19S5 (R.C.R.); Wesco Scientific Promotion Foundation (Y.S.).

## Author Contributions

R.C.R. conceived the experiments. I.C. carried out sequence analysis. I.C. and Y.S. collected and analysed the cell localization data. R.C.R. wrote the paper, with contributions from all authors.

## Author Information

The authors declare no competing financial interests.

## Materials and Methods

### Sequence analyses and structure predictions

Protein signal sequences were predicted by SignalP 6.0^13^ and cellular localization predicted by DeepLoc 2.0^14^. AlphaFold-multimer was used to predict the structures of signal peptides bound to human Sec61α^16,17^.

### Cell culture and imaging

HeLa cells were grown in Minimum Essential Media (MEM, Sigma-Aldrich) supplemented with L-glutamine and 10% fetal bovine serum (FBS) (Nichirei), and incubated at 37 °C, 5% CO_2_. Mycoplasma contamination in cell cultures was routinely tested using the PCR mycoplasma detection set (Takara Bio). At approximately 70% confluence, HeLa cells were co-transfected to express each of EGFP-fused MK-D1 proteins with an endoplasmic reticulum (ER) localizing mCherry construct using the Xfect transfection reagent (Takara Bio). At 24 h post-transfection, cells were washed with PBS (pH 7.4), fixed with 4% paraformaldehyde (Nacalai Tesque, Inc.) in PBS for 15 min at room temperature, mounted with Fluoro-KEEPER antifade reagent with DAPI (Nacalai Tesque, Inc.), and imaged using FluoView FV1200 confocal microscope (Olympus).

### Statistics and Reproducibility

The protein localization experiments were repeated twice. Pull-down and Western blots were performed twice

